# The Biogeography of Fungal Communities Across Different Chinese Wine-producing Regions Associated with Environmental Factors and Spontaneous Fermentation Performance

**DOI:** 10.1101/2020.10.22.351585

**Authors:** Ruilong Li, Siyu Yang, Mengyuan Lin, Sijiang Guo, Xiaoyu Han, Mengmeng Ren, Le Du, Yinghui Song, Yilin You, Jicheng Zhan, Weidong Huang

**Author notes:** Address correspondence to Weidong Huang,; Jicheng Zhan.

## Abstract

Chinese Marselan grapes are believed to possess the potential to become a characteristic regional variety, whose quality is internationally recognized. The fermentation-related mycobiota from six climatically diverse Marselan-producing regions in China were analyzed via high-throughput sequencing (HTS), while the influence of environmental factors was evaluated as well. The results implied that the phyla Ascomycota and genus *Aureobasidium* dominated the fungal communities in 166 Marselan must and ferment samples. Significant differences were detected in the fungal microbiota from the regions, as well as the wineries, while these discrepancies decreased as the fermentation progressed. Moreover, the difference in fungal communities between the wineries exceeded the variation involving the regions, and higher diversity was identified in the types of species than in the compositions. Geoclimatic elements (Gc) and physicochemical indexes (Pi) exerted a significant effect on the fungal must consortium, explaining 58.17% of the taxonomic information. Therefore, a correlation was proposed between the spontaneous fermentation performance, as well as the polyphenol (PP) content, and their association with fungal taxonomic composition. In addition to enriching the knowledge regarding the wine microbiome and its influencing factors, this study may provide a new strategy for harnessing autochthonous “microbial terroir”.

**Importance:** Wine microbiome and their metabolism play a crucial role in wine quality. Chinese wine-producing regions are widely distributed with diverse geoclimatic conditions, this study reports a most comprehensive biogeography of fermentation-related fungal communities performed to date, taking the Chinese promising Marselan grape variety as the research object. It reveals significant differences in the fungal microbiota of Marselan must and ferment from different regions and wineries, and higher otherness between species type than that between species composition. The study also reports the main geoclimatic and physicochemical factors shaping fungal flora. Moreover, a correlation was proposed among the spontaneous fermentation performance with fungal taxonomic composition and polyphenol content. Taken together, the results shed light on the wine fungal communities and its influencing factors, which is crucial for vineyard and fermentation microorganism management, and may also provide a new strategy for harnessing autochthonous “microbial terroir” by utilizing spontaneous fermentation.

## Introduction

The process of wine fermentation harbors a complex microbiome (1), and their metabolism plays a crucial role in wine quality (2). Although various fungi and bacteria colonize the roots, stems, leaves, flowers, and fruits of grapevines, most grape fruit-related microorganisms are unable to survive during the wine fermentation process (3), due to stress factors such as high ethanol concentrations, low pH, and anaerobic environments (4). Compared with bacteria, the fermentation process has a more substantial impact on the fungal flora (5), while the fungal must consortium displays higher annual stability (6). As the core microorganisms of alcohol fermentation, yeasts have a significant influence on the flavor and quality of wine (7), but since the microbial consortia of wine fermentation is a complex ecosystem with sophisticated interactions (8), these considerations mandate an integral understanding of fungal communities during wine fermentation (9). With the development of gene sequencing technology, several studies have documented the fungal microbiota related to wine fermentation worldwide (6, 10, 11). HTS has been widely used in fungal diversity research and has been proven an effective method for studying microorganisms during fermentation (12).

As a product with both cultural and economic value, wine is appreciated for its regional differences, also known as terroir (13). Previously, wine terroir was attributed to the soil and the vineyard environment, and the concept of “microbial terroir” was presented in conjunction with the development of microbial wine research (14). The distribution of Chinese wine-producing regions is extensive, covering 179 counties, with a range of 24-47 °N latitude (LAT) and 76-132 °E longitude (LNT). Significant differences in climate are evident between the producing regions, and various microclimates exist in each area (15). To date, research in China has focused on partial regions (16, 17), and knowledge regarding the fungal microbiome exposed to different climatic conditions nationwide remains minimal (9, 17–19).

Since being introduced into China in 2001, and due to its excellent adaptability and fermentative characteristics, the planting area of Marselan has gradually expanded. At present, China is one of the countries with the largest Marselan planting area (20). The flavor and quality of wine produced from Marselan are exceedingly popular with consumers and experts both domestically and abroad, and it is expected to become a representative variety of the Chinese wine industry (21). As an emerging variety, current studies primarily focus on its fermentation and cultivation (22), and information regarding the fungal consortium during the fermentation process of Marselan grape from different regions is scarce (23).

In this study, the fungal communities of the must and ferment from six major Marselan regions (fifteen wineries) in China are assessed via HTS analysis for internal transcribed spacer II (ITS2) genes. Furthermore, coordinate and cluster analysis are applied to test the fungal community homeomorphism among regions and wineries. In addition, correlation analysis is used to explore the relationship between the fungal consortium and geoclimatic factors, as well as the influence of fungal composition on the physicochemical indexes of spontaneously fermented wine. In conclusion, we conduct a systematic study to uncover the fungal communities and its influencing factors in different Marselan producing regions.

## Results

### Taxonomic Assignment and α-diversity Analysis of Marselan Fungal Communities

Marselan grapes from fifteen different wineries in the six main Chinese producing regions were collected, and spontaneous fermentation was carried out to investigate the fungal consortium in Marselan must and ferment. Supplementary Table S1 shows the sampling of the wineries and regions. During the fermentation process, 1611 operational taxonomic units (OTUs) with 97% similarity were observed in 166 samples with average ITS2 rDNA reads of 80,055 (Supplementary Dataset S1). Compared with the fungal reference database, taxonomic assignment revealed at least three fungal phyla (Ascomycota, Basidiomycota and Mucoromycota), 22 classes, 66 orders, 148 families, and 306 genera after removing the samples assigned to bacterial taxa, in addition to some unknown groups, indicating the relative extent of uncharacterized fungi. The Ascomycota dominated 87.68% of all OTUs, followed by Basidiomycota at 4.28% (Supplementary Dataset S2). *Aureobasidium, Alternaria*, and *Cladosporium* were the dominant fungal genera in the must samples, accounting for 24.40%, 17.29%, and 14.82%, respectively. The fungal flora of the must also contained *Colletotrichum, Rhodotorula, Metarhizium, Botrytis, Papiliotrema, Hanseniaspora*, and other trace fungi with a relative content exceeding 2%, while the unclassified genus accounted for 7.97 % (Figure. 1 and Supplementary Table S2A). Moreover, 283 fungal species were detected in the Marselan must samples of Penglai City in Shandong Province (YT), which was the highest of all six regions, while Xiangning County in Shanxi province (SX) displayed the least number at only 208 fungal species. The number of observed species in descending order was as follows: YT, Huailai County in Hebei Province (HL), Fangshan District in Beijing (FS), Changli County in Hebei Province (CL), Ningxia’s Helan Mountain’s Eastern Region (NX), and SX (Figure. 2B). Although the variations in abundance-based coverage estimator (ACE), Chao1, the Shannon indices, and the Simpson index were generally consistent with the species number trends, FS, the third-highest on the list of fungal species, exhibited the lowest Simpson index value (Supplementary Table S3A). At the winery level, Chateau Rongzi (SX.rz) in SX presented the lowest number of fungal species, while the Winery Baihuagu (HL.bhg) in HL had the highest (Figure 2E and Supplementary Table S4A). The rarefaction curves and Good’s coverage data of the Marselan must samples indicated that the ITS2 gene library was generally well-constructed (Supplementary Figure. S1A and Table S4A).

**Figure 1.**
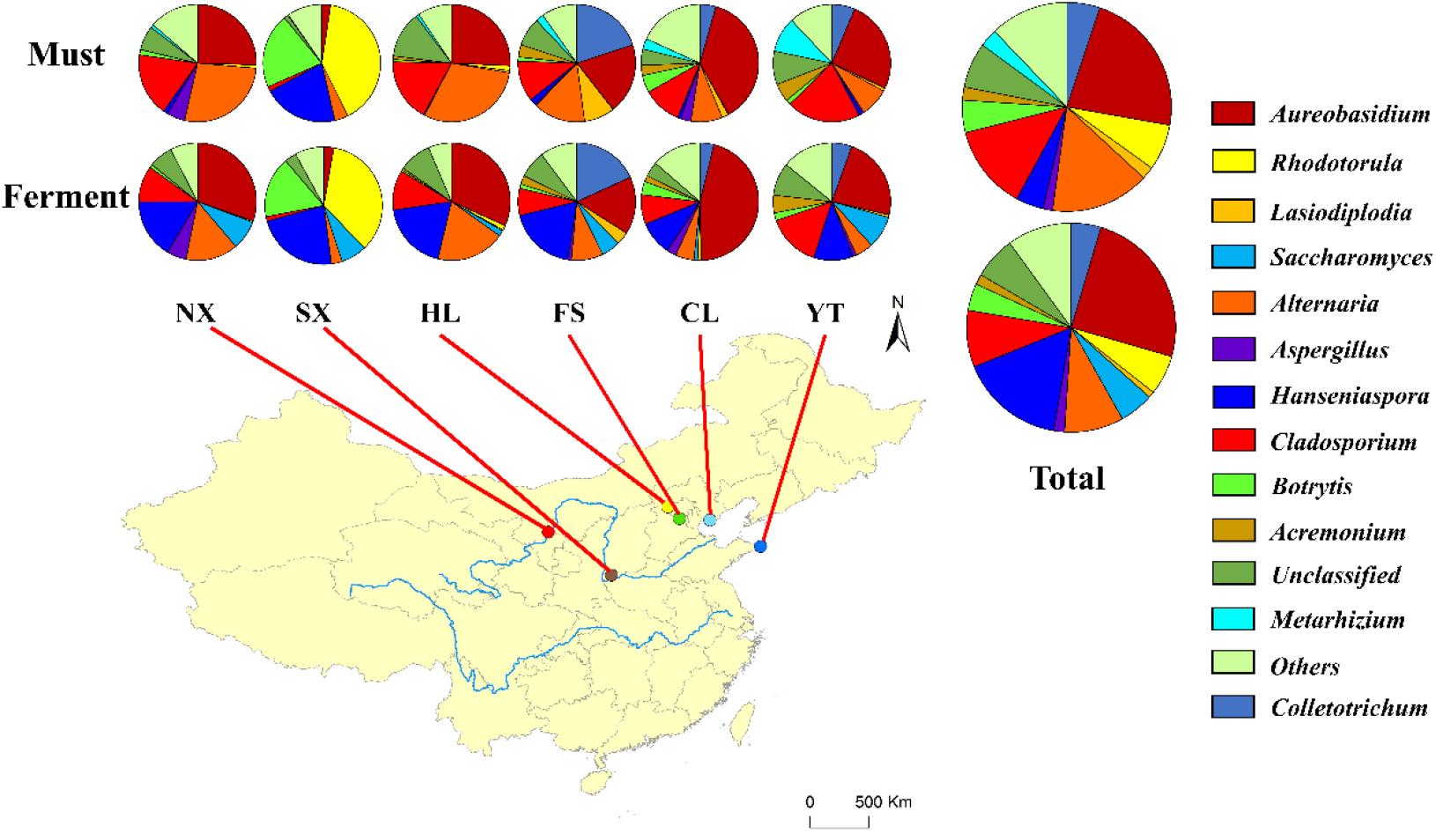
Relative abundance percentages (%) of the Marselan must and ferment fungal genera from different regions.

**Figure 2.**
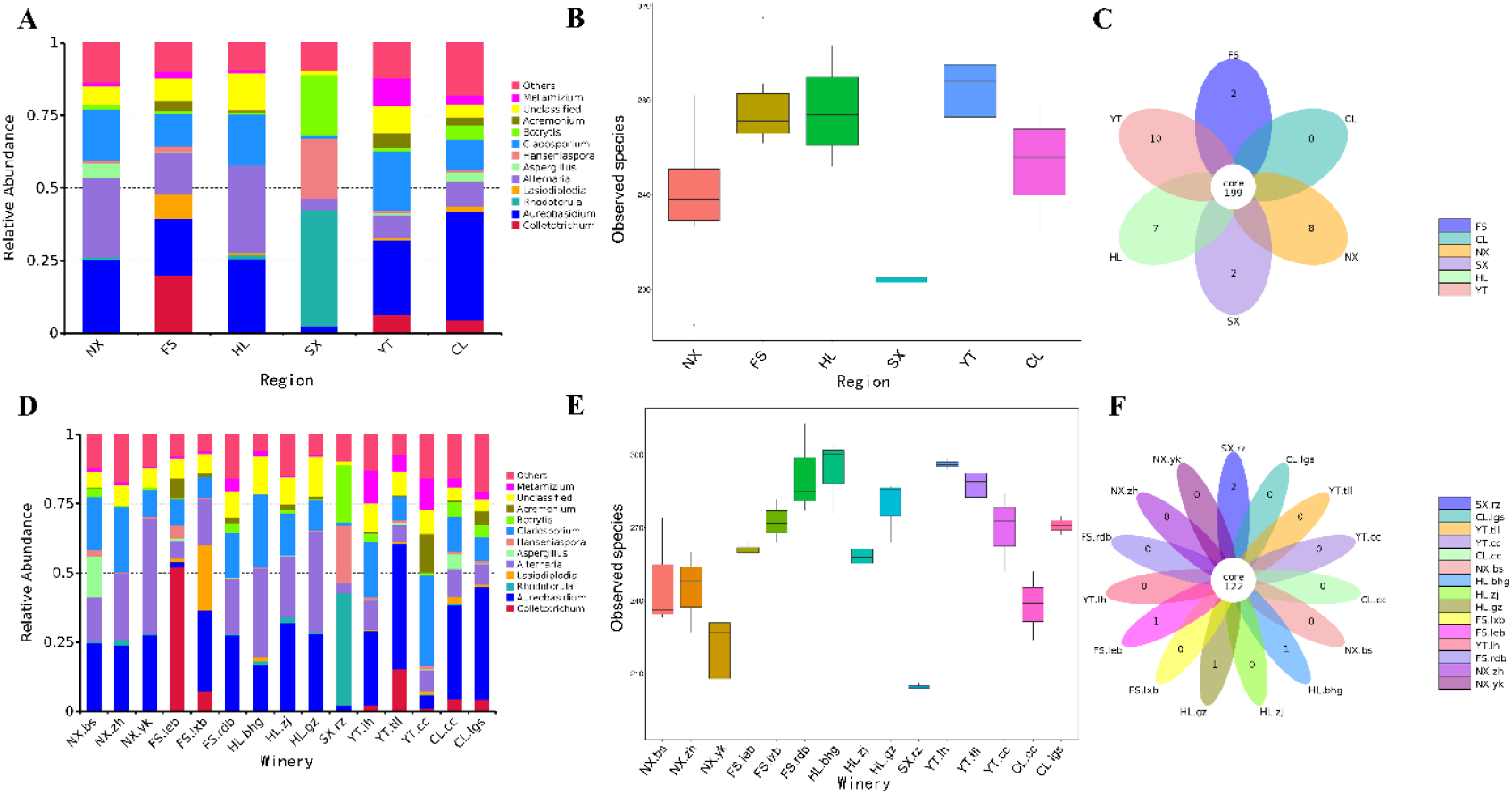
Marselan must fungal communities. (A) Relative abundance percentages (%) of the Marselan must fungal genera from different regions; (B) Wilcoxon test involving the Marselan must fungal species numbers of different regions; (C) Venn diagram of the Marselan must fungal consortium of different regions; (D) Relative abundance percentages (%) of the Marselan must fungal genera of different wineries; (E) Wilcoxon test involving the Marselan must fungal species numbers of different wineries; (F) Venn diagram of the Marselan must fungal consortium of different wineries.

Considering the fermentation process, *Aureobasidium, Hanseniaspora, Alternaria, Cladosporium*, and *Rhodotorula* comprised the primary fungal genera, accounting for 24.88%, 16.21%, 9.16%, 8.78%, and 6.14% respectively. Notably, the proportion of *Saccharomyces* throughout the fermentation process increased to 5.32%, while the unclassified genus accounted for 6.57%. The Marselan ferment fungal consortium contained *Colletotrichum*, *Botrytis*, *Metarhizium*, *Aspergillus*, *Acremonium*, and *Papiliotrema*, as well as another genus accounting for more than 1% (Figure. 1 and Supplementary Table S2B). During the fermentation process, the number of fungal species in the six Marselan regions decreased slightly, but YT still had the highest number of species and NX the lowest, while CL, ranking fourth in observed species, demonstrated the minimum Simpson index value (Supplementary Figure. S2B and Table S3B). At winery level, Winery Yunkou (NX.yk) in NX displayed the least number of species at 208, while HL.bhg in HL exhibited the highest at 299 (Supplementary Figure. S2E and Table S4B). The rarefaction curves and Good’s coverage data of the Marselan ferment samples displayed a valid ITS2 gene library (Supplementary Figure. S1B and Table S4B).

### Analysis of the Similarities and Differences between the Marselan Must and Ferment Fungal Communities of Different Regions and Wineries

Remarkably, at the winery level, only four wineries displayed unique fungal species or OTUs in the Marselan must samples, which is less than one-third of all the wineries (Figure 2F). However, at the region level, all regions contained unique fungal species except CL (Figure 2C). As a visual and intuitive display, the Non-metric multi-dimensional scaling (NMDS) method was conducted to demonstrate the similarities or differences between the fungal communities of different regions and wineries. An unweighted UniFrac NMDS plot of the Marselan must fungal community showed that although distinctions were detected among different wineries, those wineries in the same region displayed a distinct clustering phenomenon over a relatively close distance. While NX, HL, and SX were located in the left quadrant, YT, CL, and FS were located in the opposite quadrant (Figure. 3A). Considering species abundance (weighted UniFrac), the region clustering pattern still existed, although the wineries in FS and YT differed with regard to relatively extended distance, while some outliers were also evident (Figure. 3B).

**Figure 3.**
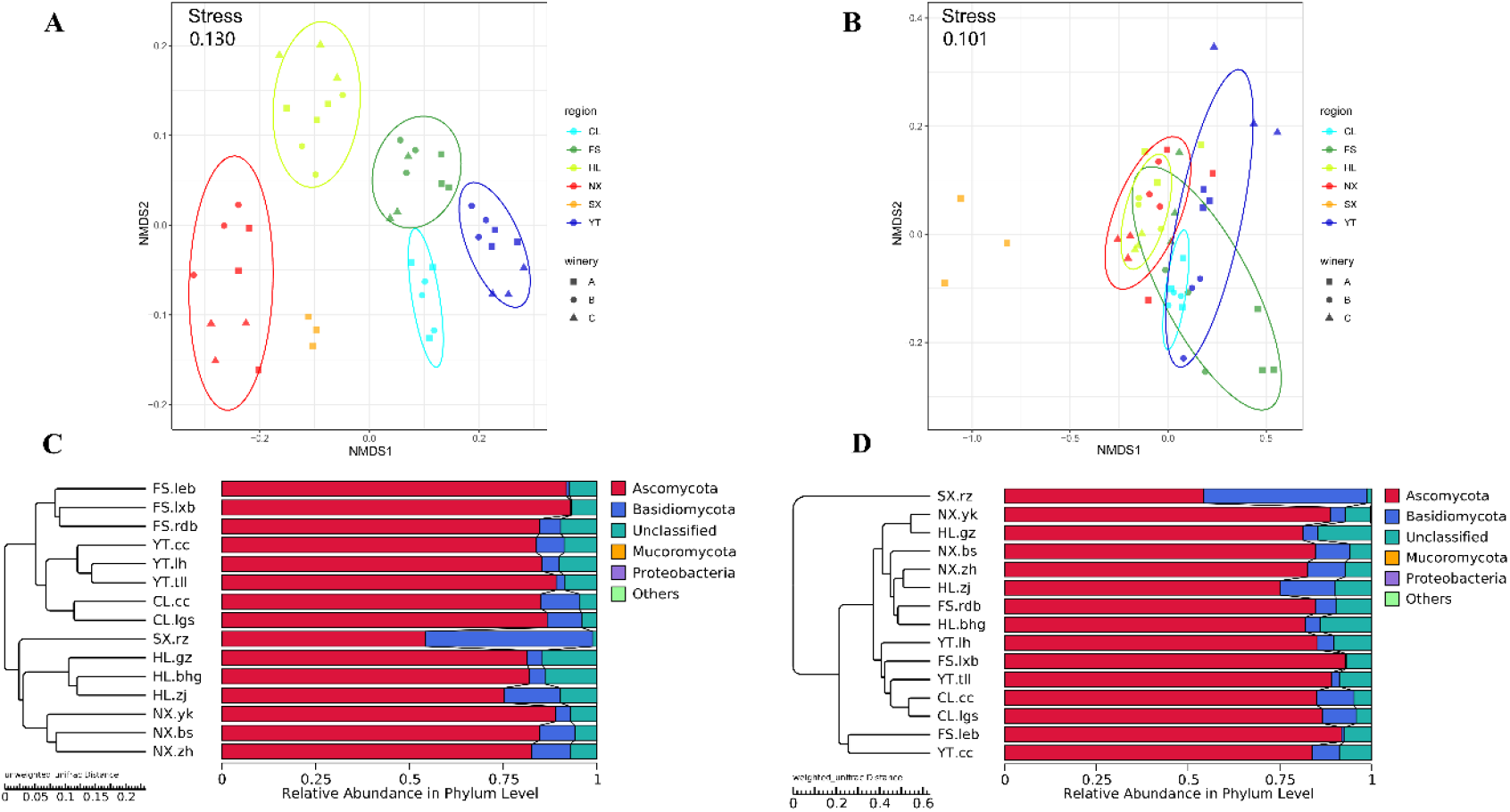
Coordinate and cluster analysis of the Marselan must fungal communities of different regions and wineries. (A) Unweighted UniFrac NMDS plot of the Marselan must fungal communities; (B) Weighted UniFrac NMDS plot of the Marselan must fungal communities; (C) UPGMA unweighted UniFrac clustering of the Marselan must fungal communities; (D) UPGMA weighted UniFrac clustering of the Marselan must fungal communities. Different colors correspond to different regions, and different shapes correspond to different wineries within a region.

The unweighted pair group method with arithmetic mean (UPGMA) clustering analysis of the Marselan must fungal communities based on the unweighted UniFrac distance further confirmed that all the wineries in the same region were clustered together. Moreover, the fungal microbiota of NX, HL, and SX was relatively similar as a branch, while the fungal communities of YT, CL, and FS regions formed the other branch (Figure. 3C). The weighted UniFrac UPGMA clustering analysis indicated that most wineries in the same regions clustered together except FS and SX, whose wineries aggregated into an independent branch with unique fungal consortiums (Figure. 3D). Except for the shorter distance between the regions in the same branch, the coordinate and cluster analysis of the Marselan ferment fungal communities were consistent with those of the must (Supplementary Figure. S3).

Analysis of similarities (ANOSIM) and multi-response permutation procedure (MRPP) test were applied to evaluate the difference in the fungal communities of Marselan must and ferment from different regions and wineries. The ANOSIM and MRPP tests, based on unweighted UniFrac indicated that substantially significant differences were observed between the fungal compositions of the must from regions (R_ANOSIM_ = 0.941, *P* =0.001; A_MRPP_ =0.333, *P* =0.001) and wineries (R_ANOSIM_ = 0.930, *P* =0.001; A_MRPP_ =0.413, *P* =0.001). Given the abundance of species, the differences between fugal compositions of the must from different regions seemed to decrease (R_ANOSIM_ = 0.435, *P* =0.001; A_MRPP_ =0.296, *P* =0.001), while those of the wineries still maintained at a relatively high level (R_ANOSIM_ = 0.734, *P* =0.001; A_MRPP_ =0.518, *P* =0.001). During the fermentation process, extremely significant differences were evident in the fungal ferment consortium of both regions and wineries. However, all differences exhibited a decline, which was either based on unweighted UniFrac (Region: R_ANOSIM_ = 0.846, *P* =0.001, A_MRPP_ =0.243, *P* =0.001; Winery: R_ANOSIM_ = 0.851, *P* =0.001, A_MRPP_ =0.295, *P* =0.001), or on the consideration of species abundance (Region: R_ANOSIM_ = 0.301, *P* =0.001, A_MRPP_ =0.183, *P* =0.001; Winery: R_ANOSIM_ = 0.451, *P* =0.001, A_MRPP_ =0.287, *P* =0.001) (Table 1).

**Table 1.** ANOSIM and MRPP tests of the Marselan must and ferment fungal communities. Significant (P <0.05); extremely significant (P <0.01).

|  |  | Unweighted Unifrac |  |  |  | Weighted Unifrac |  |  |  |
| --- | --- | --- | --- | --- | --- | --- | --- | --- | --- |
|  |  | ANOSIM |  | MRPP |  | ANOSIM |  | MRPP |  |
|  | Group | R | *P* | A | *P* | R | *P* | A | *P* |
| Must | Region | 0.941 | 0.001 | 0.333 | 0.001 | 0.435 | 0.001 | 0.296 | 0.001 |
|  | Winery | 0.930 | 0.001 | 0.413 | 0.001 | 0.734 | 0.001 | 0.518 | 0.001 |
| Ferment | Region | 0.846 | 0.001 | 0.243 | 0.001 | 0.301 | 0.001 | 0.183 | 0.001 |
|  | Winery | 0.851 | 0.001 | 0.295 | 0.001 | 0.451 | 0.001 | 0.287 | 0.001 |

Multivariate analysis of variance (Adonis) produced the same results, confirming the substantially significant differences in the Marselan must and ferment fungal communities, and showing that the differences between the fungal flora of the wineries exceeded those of the different regions (Table 2), which was consistent with the MRPP results. Differences among the wineries in each region were also investigated, indicating significant differences between the fungal flora of wineries within each region except CL. The ANOSIM of the must fungal flora of the wineries in CL showed no significant differences, either based on unweighted UniFrac (*P* =0.2) (Supplementary Table S5A) or weighted UniFrac (*P* =0.4) (Supplementary Table S5B). During the development of fermentation, the differences between the fungal communities of the wineries in the different regions decreased (Table S5). Furthermore, consistent with the results of the coordinate and cluster analysis, the ANOISM and MRPP tests demonstrated the regional pattern of the Marselan must and ferment fungal compositions, along with the extremely significant difference between the wineries in the different regions.

**Table 2.** Adonis test of the Marselan must and krausen fungal communities. Significant (*P* <0.05); extremely significant (*P* <0.01).

|  |  | **Adonis** |  |  |  |
| --- | --- | --- | --- | --- | --- |
|  |  | **Unweighted UniFrac** |  | **Weighted UniFrac** |  |
| | **Group** | $R^2$ | $P$ | $R^2$ | $P$ |
| **Must** | Region | 0.607 | 0.001 | 0.56 | 0.001 |
|  | Winery | 0.768 | 0.001 | 0.83 | 0.001 |
| **Ferment** | Region | 0.440 | 0.001 | 0.308 | 0.001 |
|  | Winery | 0.541 | 0.001 | 0.478 | 0.001 |

### Representative Fungal Species of the Marselan Regions

Linear discriminant analysis (LDA) effect size (LEfSe) algorithm with a 4.0 LDA score threshold was applied to identify the discriminative fungal taxa in the Marselan must and ferment of the different regions. During the comparison of Marselan must mycoflora, 77 fungal species were verified as being differentially abundant in six regions. Of these, Filobasidiaceae (family), *Phoma* (genus), and *Aspergillus* (genus) were significantly enriched in NX; Mycosphaerellaceae (family), *Colletotrichum* (genus), and *Lasiodiplodia* (genus) were significantly enriched in FS; *Alternaria* (genus), *Filobasidium* (genus), and *Aureobasidium sp* (species) were significantly enriched in HL; genus *Rhodotorula, Botrytis, Hanseniaspora* and *Monilinia* were significantly enriched in SX; genus *Cladosporium, Metarhizium* and *Acremonium* were significantly enriched in YT; *Aureobasidium* (genus), *Papiliotrema* (genus), and *Colletotrichum viniferum* (species) were significantly enriched in CL. Among the demonstrative fungi, genus *Rhodotorula* had the highest LDA score at 5.33, followed by genus *Aureobasidium* with an LDA score of 5.24, while genus *Alternaria* also had an LDA score above 5.00 (5.11), and the family Mycosphaerellaceae displayed the lowest score at 4.02 (Figure. 4).

**Figure 4.**
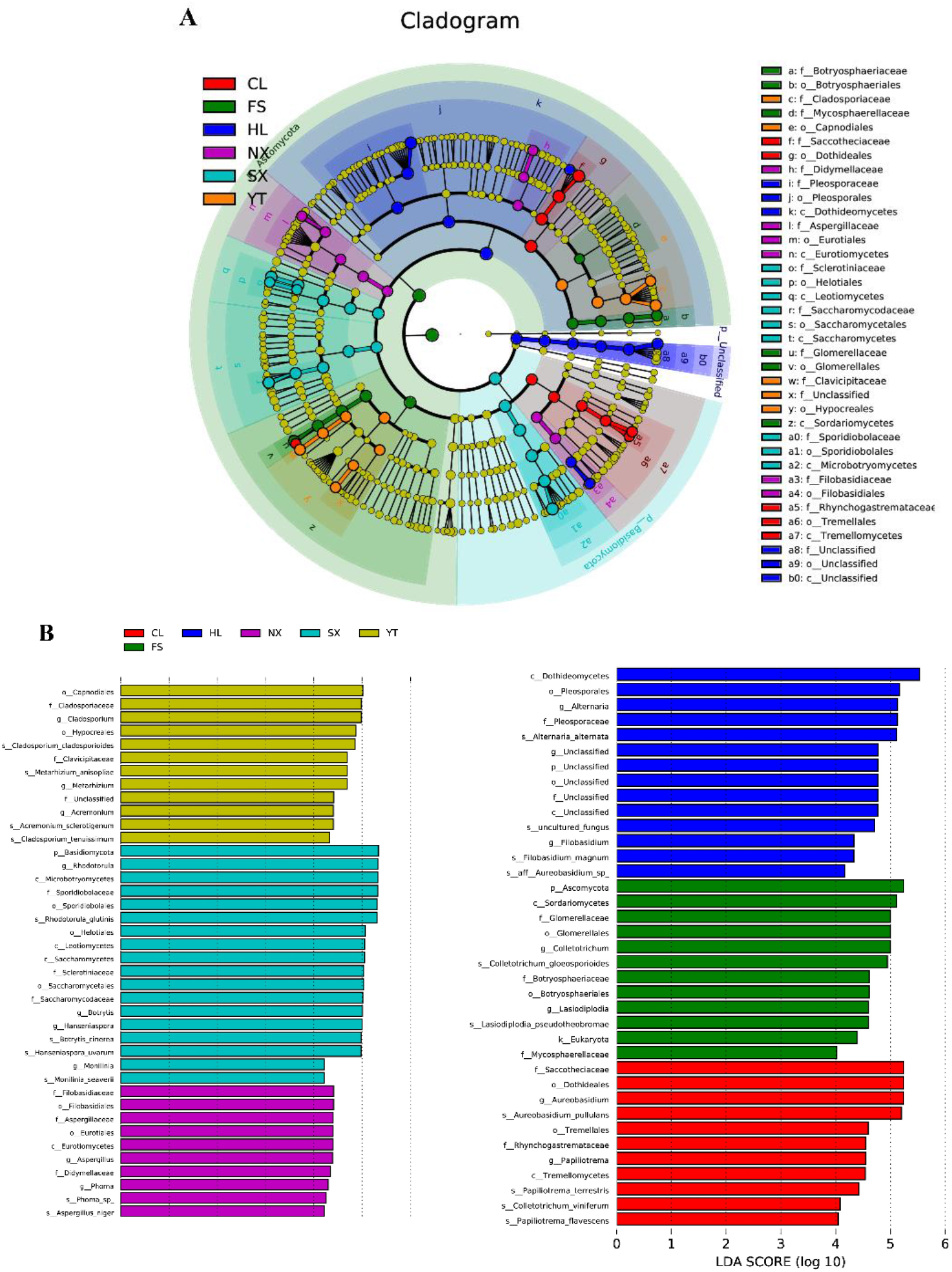
LEfSe results showing the significant fungi in the Marselan must fungal communities. (A) The cladogram reporting the taxonomic representation of the statistical and biological differences; (B) The LDA value distribution histogram.

Considering the Marselan ferment, 69 fungal taxa were verified as being differentially abundant in the six regions, which were slightly less than the Marselan must numbers. Among them, genus *Aspergillus* was the only strain exhibiting a significantly higher abundance in NX; genus *Colletotrichum*, *Lasiodiplodia*, *Candida*, and species *Hanseniaspora sp* were significantly enriched in FS; genus *Alternaria, Filobasidium* and species *Aureobasidium sp* were significantly enriched in HL; genus *Rhodotorula*, *Botrytis*, *Monilinia*, and species *Hanseniaspora uvarum* were significantly enriched in SX; genus *Cladosporium, Metarhizium*, and *Acremonium* were significantly enriched in YT; genus *Aureobasidium*, *Papiliotrema*, *Phoma*, and species *Hanseniaspora vineae* were significantly enriched in CL. Whereas, genus *Aureobasidium* had the highest LDA score of 5.35, followed by genus *Rhodotorula* with an LDA score of 5.32, while genus *Phoma* displayed the lowest score of 4.03 (Supplementary Figure S4). These results showed that of all the sampling regions used in this research, the Marselan grape regions could be clearly distinguished by the demonstrative microorganisms at different biological classification levels, from phylum to species.

### Correlation Analysis between the Marselan Must Fungal Communities and the Environmental Factors

Redundancy analysis (RDA) was applied to explore the relationship between the Marselan must fungal communities, and the environmental factors (Gc and Pi). The climatic and geographic data of 2017 for the six Marselan regions were supplied by the Wuhan Donghu Big Data Trading Center Co. Ltd. (http://www.chinadatatrading.com) of the Huafeng meteorological Media Group Co. Ltd., a subordinate unit of the China Meteorological Administration (Table 3). After removing all the correlated variables, the RDA plot indicated that ten of the 17 tested indexes significantly corresponded with the microbial community structure. These included six Gc, namely solar radiation (SR), average temperature (AT), average soil temperature (AST),LAT,LNT, and evaporation capacity (ET), as well as four Pi, namely total sugar (TS), PP, TA, and pH. While these environmental factors accounted for 58.17% of the taxonomic information, the first two axes explained 39.60% of the fungal community variation (Figure 5A). Further variation partitioning analysis (VPA) indicated that the Gc, namely SR, AT, AST, LAT, LNT, and ET were principally responsible for shaping the fungal communities of the Marselan must, accounting for 36.20% of the fungal variation, while the Pi, namely TS, PP, TA, and pH explained 10.20% of the total variability. Furthermore, 41.80% of the community distribution was not constrained by these two groups of determining factors (Figure 5B).

**Table 3.** Climatic and geographic data of the six Marselan regions in 2017 vintage.

| Region | Meteorological station | Station number | Latitude | Longitude | Average elevation (0.1m) | Average temperature (0.1°C) | Rainfall (0.1mm) | Net wind speed (0.1m/s) |
| --- | --- | --- | --- | --- | --- | --- | --- | --- |
| NX | Yinchuan City, Ningxia Province | 53614 | N38°28 | E106°12 | 11109 | 110.205 | 4226 | 15.991 |
| FS | Fangshan District, Beijing | 54596 | N39°46 | E116°12 | 489 | 133.106 | 13408 | 21.265 |
| HL | Huailai County, Hebei Province | 54405 | N40°25 | E115°30 | 5709 | 107.619 | 9498 | 24.41 |
| SX | Ji County, Shanxi Province | 53859 | N36°06 | E110°40 | 8513 | 112.695 | 12858 | 19.673 |
| YT | Fushan District, Yantai City, Shandong Province | 54764 | N37°29 | E121°14 | 539 | 135.435 | 12448 | 29.92 |
| CL | Laoting County, Hebei Province | 54539 | N39°26 | E118°53 | 85 | 132.58 | 8962 | 21.989 |
| Region | Relative humidity (1%) | Average pressure (0.1hPa) | Average soil temperature (0.1°C) | Solar radiation (0.1h) | Evapotranspiration (0.1mm) | High-temperature weather (days) | Low-temperature weather (days) |  |
| NX | 47.61 | 8910.794 | 141.849 | 28945 | 13155 | 8 | 129 |  |
| FS | 52.832 | 10111.271 | 150.504 | 25897 | 13767 | 14 | 119 |  |
| HL | 48.671 | 9502.753 | 133.498 | 31245 | 14624 | 6 | 137 |  |
| SX | 61.268 | 9197.898 | 133.09 | 21821 | 11799 | 10 | 129 |  |
| YT | 62.419 | 10107.712 | 165.479 | 25586 | 11680 | 10 | 99 |  |
| CL | 61.679 | 10160.704 | 151.213 | 26842 | 11017 | 5 | 110 |  |

**Figure 5.**
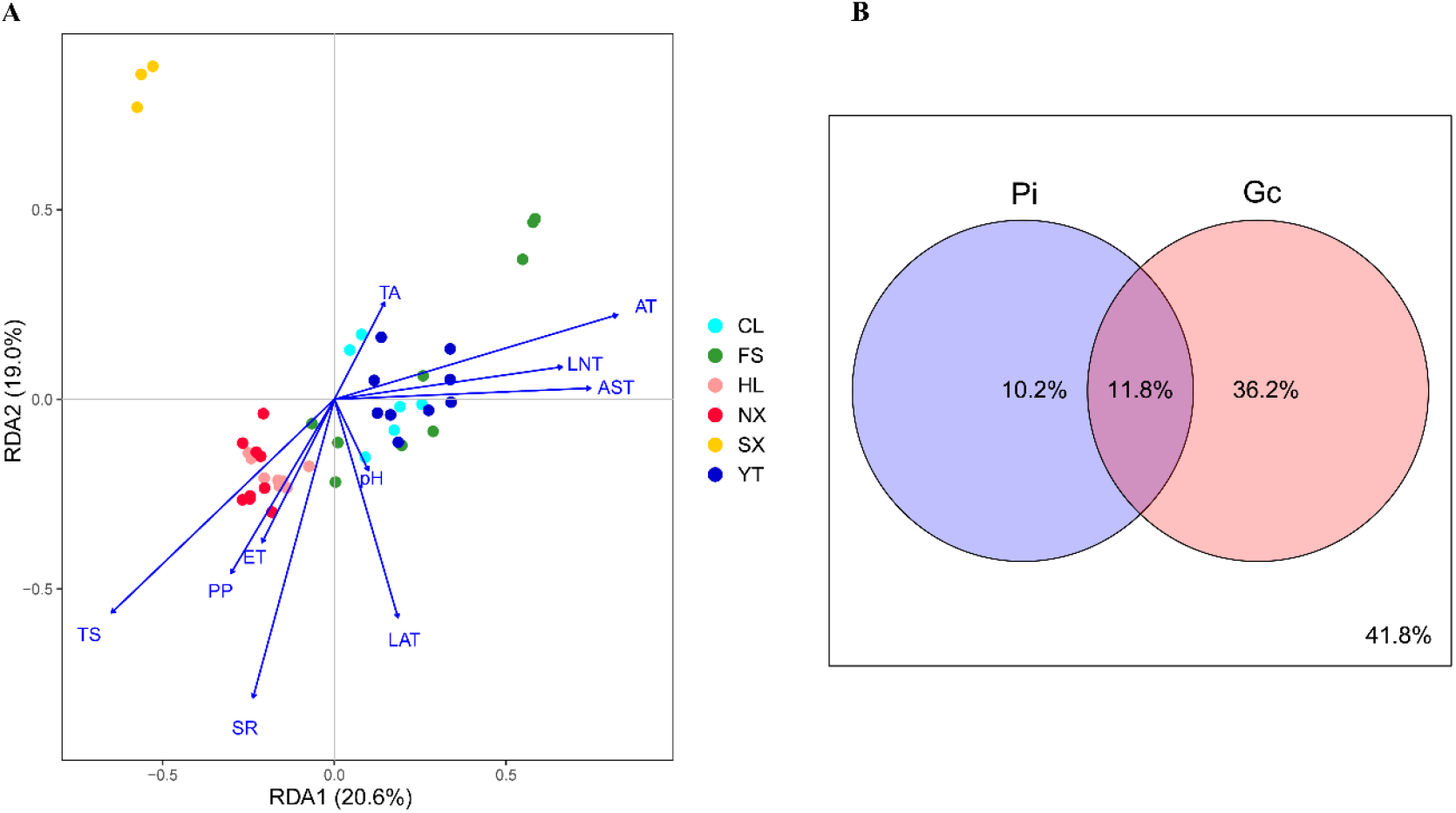
(A) RDA and (B) VPA of the environmental factors related to the Marselan must fungal communities. For the VPA, the variables presented in the RDA were separated into two groups: Gc (SR, AT, AST, LAT, LNT and ET), and Pi (TS, PP, TA, and pH).

Then, Spearman correlation analysis was used to further investigate the correlation between the variations in the major fungi in Marselan must at the genus level and the environmental factors. As such, the relative abundance of genus *Saccharomycopsis*, *Colletotrichum*, *Acremonium*, *Metarhizium*, *Lasiodiplodia* and *Penicillium* were positively correlated with LNT, AT, net wind speed (WS), relative humidity (RHD), atmospheric pressure (AP), and AST, and negatively with altitude (AE), low-temperature weather (LT) and TS. Moreover, the relative abundance of genus *Rhodotorula*, *Alternaria*, *Filobasidium* and *Naganishia* showed a significant positive correlation with AE, SR, ET, LT, and TS, and a negative correlation with LNT, AT, RHD, AP, and AST, while various some other genera of fungi were affected by the different geoclimatic, physicochemical indexes (Figure. 6).

**Figure 6.**
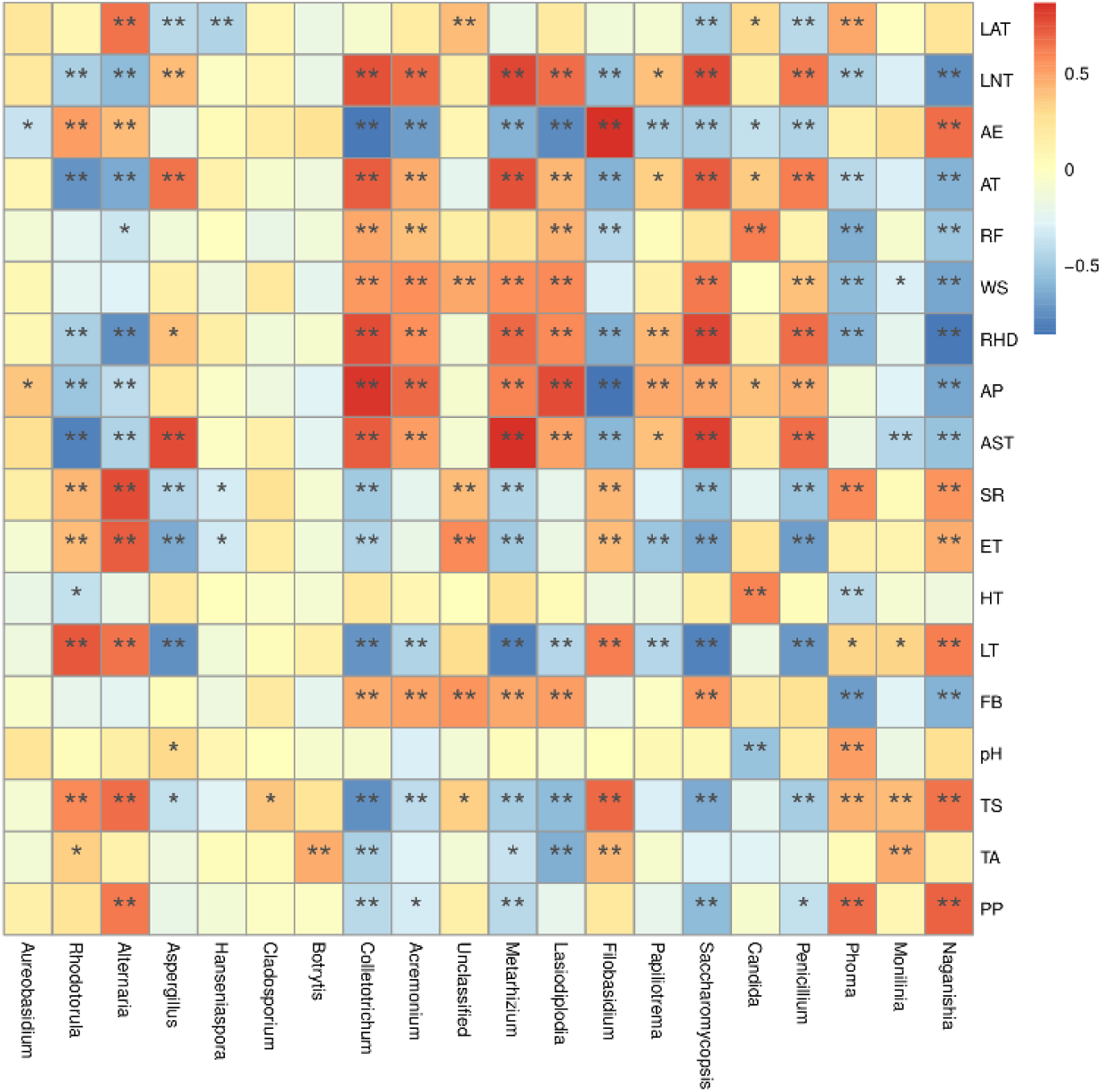
Spearman correlation analysis is used to investigate the correlation between the major fungi abundance in the Marselan must and various environmental factors. The different color intensities represent the degree of correlation. *, significant (*P* <0.05); **, extremely significant (*P* <0.01). Factors abbreviations: latitude (LAT), longitude (LNT), altitude (AE), average temperature (AT), rainfall (RF), net wind speed (WS), relative humidity (RHD), atmospheric pressure (AP), average soil temperature (AST), solar radiation (SR), evaporation capacity (ET), high-temperature weather (HT), low-temperature weather (LT), fresh breeze (FB), pH, total sugar (TS), total acid (TA), and polyphenol (PP).

### The Correlation Analysis between the Physicochemical Parameters and Ferment Fungal Communities of Spontaneously Fermented Marselan Wine

Although Marselan must generated more than 8% v/v alcohol via spontaneous fermentation (except for FS.leb, FS.rdb, HL.gz, and CL.cc), most of the wineries did not meet the residual sugar standard for dry wine of below 4g / L, including NX.zh and NX.yk in NX, FS.leb, FS.lxb, and FS.rdb in FS, HL.bhg, and HL.gz in HL, as well as CL.cc and CL.lgs in CL. Furthermore, the acetic acid (AA) content in most of the wineries exceeded the legal standard of 1.2 g/L, included NX.bs, NX.zh, and NX.yk in NX, FS.leb and FS.rdb in FS, HL.bhg in HL, as well as CL.cc and CL.lgs in CL (Table 4, OIV, 2013).

**Table 4.** The physicochemical parameters of spontaneously fermented Marselan wine. Values are given as mean ±standard deviation of three biological replicates. Letters indicate the level of significant difference (*P*<0.05) as determined with ANOVA analysis.

| Winery | Residual<br>sugars (g/L) | Ethanol<br>(% v/v) | Acetic acid<br>(g/L) | Total acid<br>(g/L) | Glycerol<br>(g/L) | pH |
| --- | --- | --- | --- | --- | --- | --- |
| NX.bs | 3.11±1.17<sup>b</sup> | 14.6±0.83<sup>a</sup> | 1.68±1.33<sup>c</sup> | 5.93±0.83<sup>b</sup> | 12.26±0.88<sup>a</sup> | 3.05±0.1<sup>c</sup> |
| NX.zh | 6.94±4.53<sup>b</sup> | 11.46±3.13<sup>ab</sup> | 1.2±1.05<sup>c</sup> | 5.43±2.15<sup>b</sup> | 9.5±1.73<sup>bc</sup> | 3.15±0.04<sup>c</sup> |
| NX.yk | 44.75±28.66<sup>ab</sup> | 8.2±5.45<sup>bcd</sup> | 5.7±5.6<sup>bc</sup> | 10.96±3.89<sup>a</sup> | 7.86±2.75<sup>d</sup> | 3.64±0.29<sup>a</sup> |
| FS.leb | 72.37±23.59<sup>a</sup> | 3.76±1.53<sup>de</sup> | 7.24±3.24<sup>b</sup> | 11.4±3.24<sup>a</sup> | 2.11±0.66<sup>e</sup> | 3.13±0.07<sup>c</sup> |
| FS.lxb | 44.48±19.28<sup>ab</sup> | 8.4±0.54<sup>bcd</sup> | 0.16±0.17<sup>c</sup> | 5.23±0.83<sup>b</sup> | 7.52±0.54<sup>d</sup> | 3.57±0.06<sup>ab</sup> |
| FS.rdb | 46.14±5.89<sup>ab</sup> | 2.55±1.8<sup>e</sup> | 1.59±1.02<sup>c</sup> | 5.43±0.85<sup>b</sup> | 3.3±1.41<sup>e</sup> | 3.45±0.04<sup>bc</sup> |
| HL.zj | 3.31±0.07<sup>b</sup> | 12.48±0.2<sup>ab</sup> | 1.25±0.26<sup>c</sup> | 4.83±0.66<sup>b</sup> | 10.85±0.81<sup>abc</sup> | 3.53±0.06<sup>ab</sup> |
| HL.bhg | 33.7±22.73<sup>ab</sup> | 10.29±1.77<sup>ab</sup> | 1.1±0.14<sup>c</sup> | 5.73±0.5<sup>b</sup> | 6.89±2.85<sup>d</sup> | 3.74±0.25<sup>a</sup> |
| HL.gz | 68.48±29.52<sup>a</sup> | 7.07±3.51<sup>cd</sup> | 1.24±0.23<sup>c</sup> | 5.3±0.6<sup>b</sup> | 6.66±1.62<sup>d</sup> | 3.77±0.11<sup>a</sup> |
| SX.rz | 0.34±0.95<sup>b</sup> | 12.86±0.54<sup>ab</sup> | 1.07±0.23<sup>c</sup> | 6±0.55<sup>b</sup> | 11.41±0.65<sup>ab</sup> | 3.31±0.15<sup>bc</sup> |
| YT.lh | 0.67±0.34<sup>b</sup> | 11.52±0.37<sup>ab</sup> | 0.91±0.46<sup>c</sup> | 5.6±0.5<sup>b</sup> | 9.04±0.08<sup>bc</sup> | 3.31±0.03<sup>bc</sup> |
| YT.tll | 1.03±0.26<sup>b</sup> | 10.66±0.18<sup>ab</sup> | 1.04±0.23<sup>c</sup> | 4.66±0.4<sup>b</sup> | 8.48±1.35<sup>bcd</sup> | 3.12±0.27<sup>bc</sup> |
| YT.cc | 4.87±7.04<sup>b</sup> | 10.28±0.6<sup>ab</sup> | 1.31±1.75<sup>c</sup> | 7.03±2.18<sup>b</sup> | 8.12±0.72<sup>bcd</sup> | 3.73±0.05<sup>a</sup> |
| CL.cc | 52.97±11.47<sup>a</sup> | 2.66±0.94<sup>e</sup> | 12.8±5.73<sup>a</sup> | 13.76±2.2<sup>a</sup> | 3.5±0.73<sup>e</sup> | 3.19±0.09<sup>c</sup> |
| CL.lgs | 13.17±10.95<sup>b</sup> | 10.76±1.76<sup>ab</sup> | 3.69±3.87<sup>bc</sup> | 9.1±4.25<sup>a</sup> | 8.23±1.64<sup>bcd</sup> | 3.23±0.02<sup>bc</sup> |

Spearman correlation analysis was preferred to investigate the correlation between the fungal compositions of the Marselan ferment and the Pi variations of the spontaneously fermented wine (difference value with must), the results indicated that anthocyanin (AC) and PP content correlated positively with genus *Filobasidium* and *Rhodotorula*, and negatively with genus *Colletotrichum* and *Acremonium*. Notably, the fermentation rate (FR) was confirmed to correlate positively with genus *Filobasidium* and *Rhodotorula*, and negatively with genus *Paramycosphaerella*, *Colletotrichum*, and *Acremonium*, which was consistent with AC and PP content (Figure. 7).

**Figure 7.**
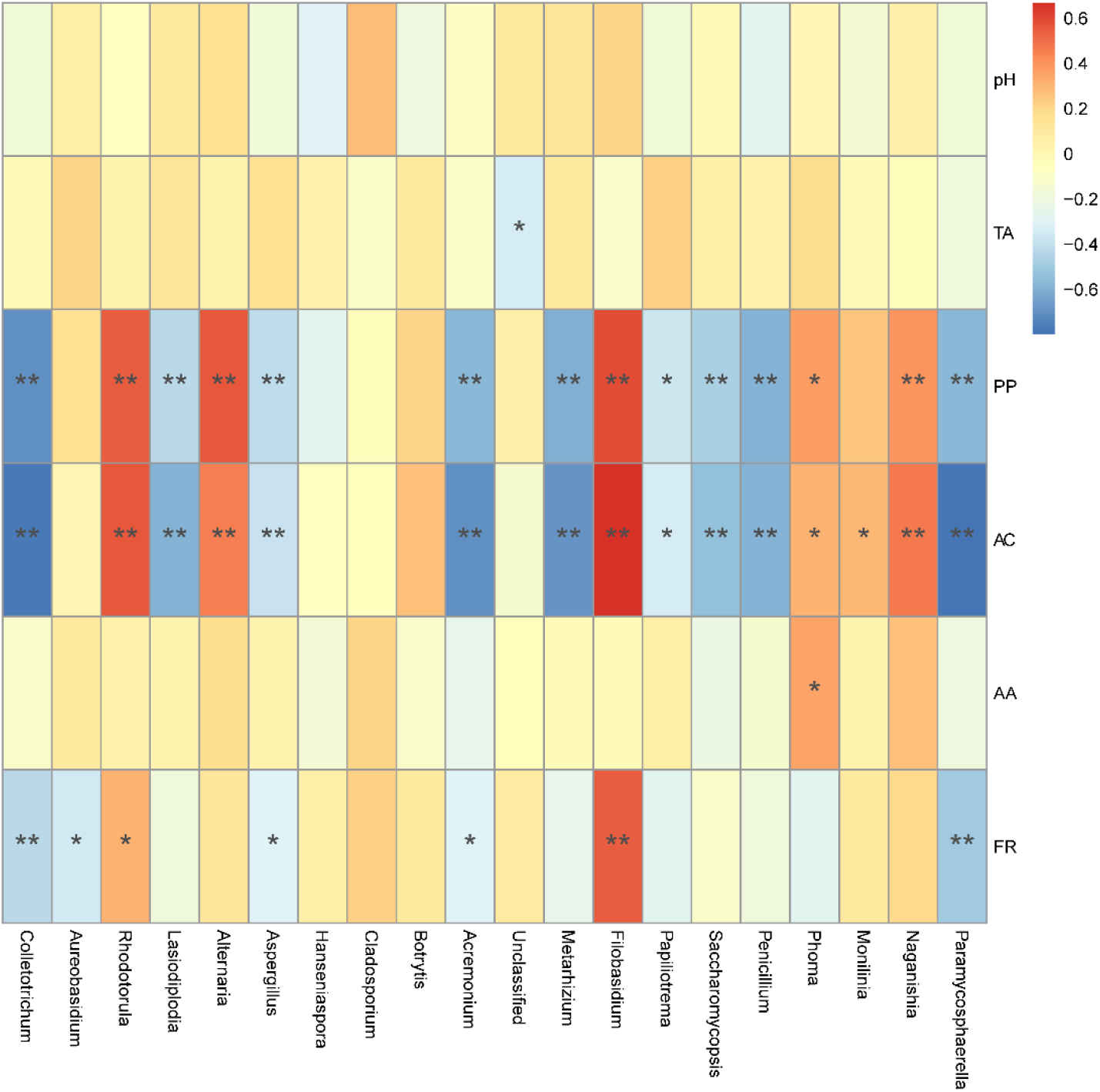
Spearman correlation analysis of the correlation between the fungal composition in the Marselan ferment and physicochemical index variations of the spontaneous fermented wine. The different color intensities represent the degree of correlation. *, significant (*P* <0.05); **, extremely significant (*P* <0.01). Factors abbreviations: total acid (TA), polyphenol (PP), anthocyanins (AC), acetic acid (AA), fermentation rate (FR).

## Discussion

The wine microbiome has been carefully studied due to its association with wine quality parameters, especially after the concept of wine “microbial terroir” was proposed (14, 24). Compared with the main grape varieties currently used for, Cabernet Sauvignon and Chardonnay, the promising Marselan variety from China has a relatively short vine age due to the late introduction (20, 23). In this study, the differences and similarities between the fungal communities in both the must and ferment of six main Chinese Marselan-producing regions, as well as the respective wineries were investigated. The correlation between fungal microbiota and environmental factors such as the Gc and Pi was also examined, further elaborating the influence of terroir conditions on fungal microbiota. This study also explored the relationship between the fungal communities and the physicochemical indexes of spontaneously fermented wine, evaluating the possibility of producing wine with regional characteristics through spontaneous fermentation by utilizing “microbial terroir.”

Despite the complexity of must microbiota and the influence of numerous factors including environmental (14) and anthropic elements (25), some common fungi always dominate. In this study, the Ascomycota and Basidiomycota still dominated at the phylum level, but the phylum Mucoromycota was also detected in a small amount, broadly corresponding with the grapevine cutting samples (26). Moreover, the frequently encountered genera *Aureobasidium*, *Aspergillus*, *Alternaria*, *Cladosporium*, *Botrytis*, *Hanseniaspora*, *Rhodotorula*, and *Filobasidiella*, were confirmed, which corresponded with similar findings involving must fungal consortiums in California, USA (6) and Spain (12). Other genera, such as *Acremonium*, and *Phoma* were also found in a vineyard in Beijing, China (19). Additionally, it is important to note that the common fungal phytopathogen genus *Colletotrichum* mainly came from winery FS.leb and YT.tll, while the proportion of this genus reached an alarming 52.2% in the FS.leb winery, and 15.3% in YT.tll (Figure 2D). Therefore, the extraordinary proportion of this pathogenic fungi caused these wineries to differ significantly from other wineries in terms of fungal composition, even when located in the same region (Supplementary Table S5). Fortunately, winery FS.leb uprooted the Marselan grapevine from the block, and planted new grape varieties after biological agents were applied.

The alcohol, high osmotic pressure, and high acid condition during the fermentation process reportedly reshaped the fungal microbiota (1), this is usually conducive to the growth of yeast (12). As the fermentation progressed, the fungal microbiota of Marselan ferment changed significantly, mainly due to the increase in the proportion of yeast and other fermentation strains, such as genus *Saccharomyces*, *Rhodotorula*, and *Hanseniaspora*, accompanied by the decrease in the proportion of other genera including *Alternaria*, *Cladosporium*, and *Metarhizium* (Figure 1). The addition of sulfur dioxide had a considerable impact on fungal population dynamics (25). However, the slight difference in the fungal compositions of the Marselan ferment may either stem from spontaneous fermentation without adding sulfur dioxide, or can be attributed to the different grape varieties and regions.

This study confirmed that the fungal microbiota of Marselan must and ferment significantly differed among the six different Chinese regions, whether species abundance was considered or not (Figure. 3, Supplementary Figure. S3, and Table 1). This result corresponded with research involving Chardonnay in USA (6) and Sauvignon Blanc in New Zealand (8). The Marselan regional pattern also broadly conformed to the conclusion that the geographical origin was more effective than the variety in impacting the fungal community (27). However, significant differences in the fungal microbiota were also detected regardless of whether the wineries were in different regions or the same region (Figure. 3, Supplementary Figure. S3, Table 1, and Supplementary Table S5). In addition, the differences between wineries seemed to be more substantial than those between regions (Table 1 and Table 2). Various factors may alter the fungal consortium intra-region, such as microclimate, soil conditions, or agricultural management (28), thus forming the unique fungal flora between different wineries. Furthermore, the differences in the fungal communities of different wineries in the same regions also reduced the variations in disparate regions. The topography of the sampling wineries in FS and YT is quite different. The FS.rdb winery is relatively far away from the other two wineries in FS and located on a slope, while the FS.leb and FS.lxb wineries are located on flat ground. Winery YT.lh is located on a slope, while wineries YT.tll and YT.cc are located on flat ground. These discrepancies may further alter the fungal compositions of different wineries within the same region, leading to a more substantial difference between the fungal compositions of the wineries in these two regions (Supplementary Table S5). Therefore, further studies are necessary to determine the influencing factors of fungal communities on blocks with different microclimatic, viticultural, and geophysical conditions beyond the scope of the measurements of this study.

Moreover, this research revealed that the otherness of the Marselan fungal consortium between species type exceeded the differences between species composition, except for the divergence between the must fungal communities of different wineries (Table 1 and Table 2). Although the wine fermentation process possesses complex fungal communities, the must fungal consortium is generally dominated by phylum Ascomycota and Basidiomycota, while the genera *Aureobasidium*, *Hanseniaspora*, *Aspergillus*, *Alternaria*, *Cladosporium*, *Candida* and *Rhodotorula* are frequently detected in comparatively high proportions (4). The demonstrative fungi distinguishing different regions are usually the genera with low abundances, such as *Penicillium*, *Colletotrichum*, *Botrytis* (27), consequently resulting in more significant differences between the fungal microbiota based on unweighted UniFrac distance than those related to weighted UniFrac distance. The illusion displayed by Wayne diagram that the microbial types were more distinct between regions than between wineries (Figure. 2 and Supplementary Figure. S2) may be due to the similarity between the microbial types from different wineries in the same region (29). It can also be ascribed to the fact that fungal community discrepancies depend on both region-specific species and the proportion of shared species (30). Furthermore, this study discovered that the differences among the Marselan fungal communities decreased during fermentation compared to the must (Table 1 and Table 2), which corresponds with reports that the Cabernet microbial community profiles became less distinct as the microbial signatures among the AVAs and vineyards diminished during fermentation (24).

Recent investigations have characterized many factors affecting the must and fermentation microbiome, including locally managed (31) and native ecosystems (27). The results indicated that Gc, such as SR, AT, AST, LAT, LNT and ET exerted a significant effect on fungal communities, revealing that approximately the same geographic and climatic factors regulated fungal microbiota (6, 16). The results further showed that Pi, such as TS, PP, TA, and pH, also modified the fermentation microflora (Figure. 5A and Figure. 6). Although the physical and chemical elements of must are closely related to geoclimatic conditions, this correlation somehow guides harvest time selection and fermentation microorganism management. However, the effect of PP on fungal consortiums must be highlighted. Since PP is not only an essential constituent of red wine (32), but is also a functional component beneficial to health (33), as well as a broad-spectrum antibacterial agent (34), further research is necessary to clarify the interaction between PP and fermentation microorganisms. As far as is known, no previous reports exist regarding the relationship between PP and fermentation fungal communities. In addition, the factors involved in the study only explained 58.17% of the taxonomic information (Figure. 5B), with 41.83% unknown influence confirming the other factors affecting fungal communities (31, 35).

Spontaneous fermentation is an effective measure to maximize the role of autochthonous “microbial terroir” (11). This study detected the quality-related physicochemical indicators of naturally fermented wine, such as TS, TA, AA, and ethanol, to assess the feasibility of spontaneous fermentation. Although the naturally fermented wine from many of the wineries did not meet the dry wine standard set by the OIV (Table 2), the results indicated a positive correlation between FR and genera *Filobasidium* and *Rhodotorula*, while negatively correlating with *Colletotrichum* and *Acremonium*. It is worth mentioning that these connections to fungal communities also applied to AC and PP content (Figure. 7). With increasing evidence supporting the role of microbial terroir in shaping regional wine phenotypes (24, 36), it may provide new approaches for harnessing spontaneous fermentation to produce wines with regional characteristics by improving the content of relevant grape components beneficial to this type of fermentation process.

## Materials and Methods

### Grape Sampling and Laboratory-scale Spontaneous Fermentation

The grape samples were collected from six Marselan producing regions in China, namely, NX, FS, HL, SX, YT and CL in 2017 vintage. The sampling sites included the Jing-Jin-Ji Region (FS, HL, CL) and Loess Plateau Region (SX), all of which had warm climates, and were characterized as semi-humid zones; the Shandong Region (YT) with a hot climate, was regarded as a semi-humid zone; and the Helan Mountain East Region (NX) with a temperate climate which was classified as an arid zone, according to the frost-free season (FFS) and dryness index (DI). Furthermore, the grapevines in the SX and YT could overwinter without soil burial, but the grapevines in FS, HL, CL and NX had to be buried with soil for protection against the winter chill (15). For each region, three of the most representative wineries were considered for sampling, except for SX and CL, in which only one and two wineries, respectively, had planted Marselan grapevines (Supplementary Table S1).

For all regions, the Marselan grape samples were picked on a sunny morning within 3 d of harvest. Private wine producers authorized the sampling, and the field study did not involve any endangered or protected species. For each winery, 2 kg of healthy and undamaged grapes were collected using garden scissors sterilized with 75% alcohol. To ensure the representativeness of the sampling, multiple bunches of grape samples were randomly picked from different positions in the vineyards, containing different ranks and orientations. These samples were placed into sterile plastic bags and transported to the laboratory chilled on ice within 1 d (5).

Here, 45 grape samples from fifteen wineries (three parallel samples per winery) were aseptically destemmed, and hand-squeezed in a clean bench, after which1050 mL of grape must and pomace were equally divided and placed into three sterile 500 mL jars and sealed with sterile sealing films. The physical and chemical indexes of the initial must is shown in Supplementary Table S8. Stationary spontaneous fermentation was performed in duplicate at room temperature (controlled at 25 ± 2°C) (37).

According to the pre-experiment of 2016 and the fermentation process of 2017, the microbial diversity of the Marselan must, and krausen samples were analyzed in four stages: the must, corresponding to the juice of the crushed grapes; the start of alcoholic fermentation, which corresponded to day 3 when a rapid reduction in the °Brix level was evident; the mid-stage of alcoholic fermentation, which corresponded to day 5 when the °Brix had declined by a third; and the end of alcoholic fermentation, which corresponded to day 8 when the °Brix level was reduced by half. It should be noted that the ferment samples contained samples from all the four stages, including the must samples, and krausen samples from the start, mid-stage, and end of alcoholic fermentation. During each stage, 9 mL must, or krausen samples were collected, placed into two sterile 10 mL centrifuge tubes (one set as a backup sample), and stored at −80 °C for DNA extraction. In addition, during the start of alcoholic fermentation sampling, the fermented jars were sealed with sterile plastic and sealing films to provide anaerobic fermentation conditions. During the mid-stage of the alcoholic fermentation sampling, the fermented mash was transferred from the 500 mL jars to 250 mL sterilized triangular flasks to remove the skin residue. The fermentation process was monitored by measuring the °Brix every 2 d, and fermentation was considered complete when the °Brix displayed no change three consecutive times, and the clear ferment on day 20 was taken as the naturally fermented wine sample. Then, respective 50 mL must, and naturally fermented wine samples were stored at −20 °C for physical and chemical index analysis. The fermentation process is shown in Supplementary Figure. S5.

### DNA Extraction, PCR Amplification and ITS2 rDNA Sequencing

The genomic DNA of the fungal communities in the Marselan must, and ferment samples were extracted using a modified CTAB method (38). The purity and concentration of the DNA were quantified with a nano spectrophotometer (Thermo Scientific, Wilmington, DE, USA) and 1% (w/v) agarose gel. The fungal ITS2 gene PCR was performed via the ITS3-2024F (5’-GCATCGATGAAGAACGCAGC −3’) and ITS4-2409R (5’-TCCTCCGCTTATTGATATGC −3’) primers. All the samples were amplified in triplicate, and no-template controls were included in all steps of the process. The Qiagen Gel Extraction Kit and PCR Clean-up (Qiagen, Germany) were employed to refine the PCR products. Furthermore, fluorometric evaluation using the Qubit 2.0 dsDNA HS Assay Kit (Invitrogen, Carlsbad, CA, USA) obtained the purity of the PCR mixture, after which sequencing analysis continued via the IonS5TMXL sequencing platform (IonS5TMXL, Thermofisher, USA) from Novogene, Beijing, China.

### Measurement of Physicochemical Parameters

The pH and °Brix of the samples were measured with a HANNA 211 pH meter (HANNA, Padova, Italy) and a refractometer (ATAGO, Tokyo, Japan), respectively.

The glucose, fructose, ethanol, and glycerol content were determined with HPLC using a Waters 2414 RI Detector and a BIO-RAD Aminex HPX-87H resin-based column (300*7.8 mm) (39), and eluted with 5 mM H2SO4 at 55 °C, 0.5 mL/min.

The AA, TA, PP and AC were analyzed with corresponding Randox kits on the Randox Monaco Analyzer (Randox, Monaco, UK).

### Data Analysis

Raw sequencing reads obtained from the IonS5TMXL platform were paired and pre-merged using FLASH software (Version 1.2.7), as well as filtered with the QIIME software (Version 1.7). All quality filtered sequencing reads were then clustered into OTUs with a minimum identity of 97%, by applying UPARSE software (Version 7.0) (40). Additionally, the OTU table underwent a series of filtering steps, including removing low-quality bases and chimeras, removing the OTUs with < 3 counts across all samples, removing possible contaminants (mitochondrial and chloroplast sequences). For each fungal representative sequence (OTUs), taxonomy was assigned based on the UNITE fungal ITS database. Based on the Shannon index, the replicates were examined for outliers, resulting in the removal of fourteen samples. The remaining OTUs samples were rarefied at a value equal to the median amount of sequences (80160 sequences) to compensate for the uneven sequencing depth between the samples.

The Shannon index, Simpson index, ACE, chao1, and phylogenetic diversity (PD) whole tree values were calculated to compare the intra-group diversity (alpha diversity). A Venn diagram was applied to display the number of shared and unique OTUs in different wineries and regions (41). Moreover, Wilcoxon tests were used to evaluate the differences in average alpha diversity indices between the groups (42). NMDS was applied based on both qualitative (unweighted UniFrac) and quantitative (weighted UniFrac) distance metrics to illustrate the similarities between the wineries and the regions, using QIIME, the phyloseq (version 1.24.0) and vegan (version 2.5.6) R package (43). The UPGMA were used based on the unweighted and weighted UniFrac distances to verify the clustering of the different groups (44). Moreover, the ANOSIM, ADONIS and the MRPP was conducted to identify the significant differences between the wineries and the regions using the vegan R package (45).

Subsequently, the linear LEfSe algorithm was performed to identify the representative fungal taxa of each region, by utilizing the Huttenhower Lab Galaxy Server (40). After the environmental factors (Gc and Pi) were screened via the variance inflation factor (VIF) and step selection using the vegan and dplyr (version 0.4.3) R package, detrended correspondence analysis (DCA) result determined to apply RDA to explore the correlation between environmental factors and fungal flora. Moreover, VPA was performed to evaluate the contribution rate of each factor (46). In addition, Spearman’s correlation analysis was applied to estimate the correlation between the environmental factors, the Pi variations of the spontaneously fermented wine (difference value with must), and the major fungi genus (47).

Other data were expressed as mean ± SD, and the statistical significance between the groups was analyzed with one-way analysis of variance (ANOVA) using SPSS 19.0 software (SPSS Inc., Chicago, IL, USA).

### Data Availability

The sequencing data have been deposited in the NCBI database (https://www.ncbi.nlm.nih.gov/) under the accession number SRP269142.

## Acknowledgments

This research was supported by the National Key R&D Program of China, grant number: 2016YFD0400504. We would like to thank Qian Yu in Chateau Changyu Mosel XV, Jinhua Zhou in Chateau Huaxia Greatwall, Leipeng Wei in Chateau Rongzi, Zhe Yang in Chateau Longxi, Jianhua Luo in Guizu Winery, and Zhu Wang in Amethyard for arranging to collect samples.

## Author Contributions

Conceptualization, Ruilong Li, Jicheng Zhan and Weidong Huang; Data curation, Ruilong Li, Siyu Yang and Mengyuan Lin; Formal analysis, Ruilong Li; Funding acquisition, Weidong Huang; Investigation, Ruilong Li, Siyu Yang, Sijiang Guo, Mengyuan Lin and Xiaoyu Han; Resources, Siyu Yang, Le Du and Yinghui Song; Supervision, Jicheng Zhan and Weidong Huang; Writing – original draft, Ruilong Li; Writing – review & editing, Xiaoyu Han, Mengmeng Ren and Yilin You. All authors have read and agreed to the published version of the manuscript.

## Supplementary Materials

**Figure S1** Rarefaction curves of Marselan must (A) and krausen (B) samples.

**Figure S2** Marselan krausen fungal communities.

(A) Relative abundance percentages (%) of the Marselan krausen fungal genera from different regions; (B) Wilcox test involving the Marselan krausen fungal species numbers of different regions; (C) Venn diagram of the Marselan krausen fungal consortium of different regions; (D) Relative abundance percentages (%) of the Marselan krausen fungal genera of different wineries; (E) Wilcox test involving the Marselan krausen fungal species numbers of different wineries; (F) Venn diagram of the Marselan krausen fungal consortium of different wineries.

**Figure S3** Coordinate and cluster analysis of the Marselan krausen fungal communities of different regions and wineries.

(A) Unweighted UniFrac NMDS plot of the Marselan krausen fungal communities; (B) Weighted UniFrac NMDS plot of the Marselan krausen fungal communities; (C) UPGMA unweighted UniFrac clustering of the Marselan krausen fungal communities; (D) UPGMA weighted UniFrac clustering of the Marselan krausen fungal communities. Different colors correspond to different regions, and different shapes correspond to different wineries within a region.

**Figure S4** LEfSe results showing the significant fungi in the Marselan krausen fungal communities.

(A) The cladogram reporting the taxonomic representation of the statistical and biological differences.; (B) The LDA value distribution histogram.

**Figure S5** Spontaneous fermentation process of Marselan grape.

(A) Experiment process; (B) The variation of °Brix.

**Table S1** Sampling regions and wineries of Marselan grape.

**Table S2** Relative proportion of Marselan fungal must (A) and krausen (B) consortium.

**Table S3** Fungal α-diversity of Marselan must (A) and krausen (B) from different regions based on internal transcribed spacer II (ITS2) of rDNA analysis.

**Table S4** Fungal α-diversity of Marselan must (A) and krausen (B) from different wineries based on internal transcribed spacer II (ITS2) of rDNA analysis.

**Table S5** ANOSIM, MRPP, and Adonis tests of the Marselan must and krausen fungal communities in different wineries of different regions based on unweighted UniFrac (A) and weighted UniFrac (B) distance. Significant (*P* <0.05); extremely significant (*P* <0.01).

**Table S6** Physical and chemical indexes of the Marselan must in different wineries.

**Database S1** OTUs detected in the 166 Marselan must and ferment samples

**Database S2** Taxonomic assignment and species relative proportion of the 166 Marselan must and ferment samples.

